# Ghrelin signalling in AgRP neurons links metabolic state to the sensory regulation of AgRP neural activity

**DOI:** 10.1101/2023.05.28.542625

**Authors:** Wang Lok So, Jiachen Hu, Lotus Jeffs, Harry Dempsey, Sarah H. Lockie, Jeffrey M Zigman, Romana Stark, Alex Reichenbach, Zane B. Andrews

## Abstract

**Objective:** The sensory detection of food and food cues suppresses Agouti related peptide (AgRP) neuronal activity prior to consumption with greatest suppression in response to high caloric food or energy need. Although external sensory cues regulate AgRP neuronal activity, the interoceptive mechanisms priming an appropriate AgRP neural response to sensory information of caloric availability remain unexplored. Since hunger increases plasma ghrelin, we hypothesized that ghrelin receptor (GHSR) signalling on AgRP neurons is a key interoceptive mechanism integrating energy need with external sensory cues predicting caloric availability.

**Methods:** We used in vivo photometry to measure the effects of ghrelin administration or fasting on AgRP neural activity with GCaMP6s and dopamine release in the nucleus accumbens with GRAB-DA in mice lacking ghrelin receptors in AgRP neurons.

**Results:** The deletion of GHSR on AgRP neurons prevented ghrelin-induced food intake, motivation and AgRP activity. The presentation of food (peanut butter pellet) or a wooden dowel suppressed AgRP activity in fasted WT but not mice lacking GHSRs in AgRP neurons. Similarly, peanut butter and a wooden dowel increased dopamine release in the nucleus accumbens after ip ghrelin injection in WT but not mice lacking GHSRs in AgRP neurons. No difference in dopamine release was observed in fasted mice. Finally, ip ghrelin administration did not directly increase dopamine neural activity in the ventral tegmental area.

**Conclusions:** Our results suggest that AgRP GHSRs integrate an interoceptive state of energy need with external sensory information to produce an optimal change in AgRP neural activity. Thus, ghrelin signalling on AgRP neurons is more than just a feedback signal to increase AgRP activity during hunger.

## 1. Introduction

The brain maintains energy homeostasis by balancing energy intake with expenditure through a large neural network primarily involving the hypothalamus and brainstem. However, more recent studies suggest that the neural network governing food intake and energy balance extends beyond these regions to integrate with cortical, subcortical and limbic structures [1; 2,Clarke, 2023 #5667; 3]. Within the hypothalamus, agouti-related peptide (AgRP) neurons in the arcuate nucleus (ARC) are a critical population responsible for increasing food intake and preventing the adverse consequences of starvation [4; 5]. AgRP neurons are sensitive to interoceptive hunger-sensing cues, since photostimulation of AgRP neurons in ad libitum mice drives a learned operant sequence previously rewarded only in a fasted state [6]. Importantly, photostimulation of gamma-aminobutyric acid (GABA) neurons in the lateral hypothalamus (LH; LH^GABA^ neurons), which increases food intake [7], do not show the same hunger sensing properties [6]. Thus, although other neurons, such as LH^GABA^ neurons increase food intake, AgRP neurons are considered the canonical drivers of food intake and metabolism in response to energy deficit and homeostatic need. Further evidence for this conclusion includes the increased electrophysical firing properties, transcriptional activity, spine formation and peptide release of AgRP neurons during hunger [8–11]. Moreover, AgRP neurons are required for an appropriate feeding response to cold exposure to meet increased thermogenic energy demands [12], highlighting the importance of AgRP neurons to homeostatic need rather than just energy deprivation caused by fasting. Artificial AgRP neuronal activation mimics the effects of energy deficit and homeostatic need by enhancing food intake and reducing energy expenditure [13–16]. Their important role in energy balance is underscored by the body weight loss and starvation associated with genetic ablation in adulthood [17–19]. [20; 21]

The control of energy homeostasis by neuroendocrine feedback dictates that hunger increases AgRP activity, in response to a peripheral energy imbalance, thereby increasing food intake and decreasing expenditure to restore energy balance. Important feedback signals associated with hunger include high plasma ghrelin, which increases AgRP activity [22; 23], and low leptin and insulin, which lowers the inhibitory action of these hormones on AgRP neurons [24; 25]. However, the discovery that AgRP neurons are inhibited by sensory cues of food availability and caloric density [9; 26; 27] highlights an important, yet underappreciated, role of synaptic input on AgRP activity and energy homeostasis. These studies show that food presentation or learned food cues cause a rapid inhibition of AgRP activity (0-10 secs) prior to any meaningful calorie consumption. The fall in AgRP activity is scaled to caloric availability with peanut butter or caloric gels causing a greater suppression compared to chow diet or caloric-free gels [27; 28]. This sensory input has been traced to an afferent inhibitory input from GABA neurons in the dorsomedial hypothalamic nucleus (DMH) [29]. Gut peptide signalling and gut-brain feedback is required to confirm sufficient calorie consumption maintain AgRP activity at low levels [28; 30; 31]. Thus, an updated model of energy homeostasis should integrate mechanisms controlling rapid sensory input, as well as negative feedback, on neural circuits controlling energy homeostasis.

In addition to caloric density affecting the fall in AgRP activity, an animal’s internal energy need is also equally important since the fall in AgRP activity to the same food is greater in fasted compared to fed mice [27; 32]. This observation suggests that AgRP neurons integrate internal cues of homeostatic energy need together with external sensory cues predicting food availability. In many circumstances the balance between internal energy needs and food availability in an external environment can influence behaviour. For example, pre-emptive AgRP neuronal stimulation drives fed mice to seek food reward despite the threat of shock [33]. Therefore, interoceptive mechanisms of energy need influence the salience of sensory inputs onto AgRP neurons.

While numerous studies have assessed the nature of the synaptic input onto AgRP neurons [29; 34] or the gut-brain feedback coupled to calorie consumption [28; 30; 31], the interoceptive mechanisms priming AgRP neuronal responses to fasting remain unknown. We hypothesize that such a mechanism should involve negative feedback conveying energy need prior to the sensory detection of food or learned food cues. One potential feedback mechanism is an increase in plasma ghrelin during fasting [35] acting through ghrelin receptors (Growth Hormone Secretagogue Receptor; GHSR) on AgRP neurons (AgRP^GHSR^). In many cases the actions of ghrelin phenocopy those of AgRP neurons; 1) ghrelin is a bona fide hunger signal as ghrelin injection to ad libitum fed mice elicits an operant behavioural response learned during hunger (fasting) [6; 36]; 2) ghrelin increases food intake, motivation and dopamine neural circuits [22; 37-39]; 3) ghrelin supresses energy expenditure and lipid utilisation [40; 41]; and 4) ghrelin reduces anxiety-like behaviour and increases exploratory behaviour [42–44]. All of the above functions described for ghrelin and/or GHSR signalling have also been linked to AgRP neurons [16; 32; 45-48] and indeed some show a direct role of GHSR signalling in AgRP neurons on food intake, meal duration and energy expenditure and thermogenesis [49–51]. In this study, we hypothesized that GHSR expression on AgRP neurons is a key interoceptive mechanism integrating energy need with the salience of external sensory detection of food or food cues. To do this, we examined whether GHSR deletion in AgRP neurons affected AgRP neural activity or dopamine release in the nucleus accumbens (NAc) using *in vivo* fibre photometry in response to food or non-food objects.

## Methods

### 1.1 Mice and housing

Mouse experiments were conducted in compliance with the Monash University Animal Ethics Committee guidelines. External environment was maintained under standard laboratory conditions at 23°C in a 12-hour light/dark cycle with *ad-libitum* access to chow (chow diet, catalog no. 8720610, Barastoc Stockfeeds, Victoria, Australia) and water. Male mice 8 weeks or older were used for experimentation and group-housed unless destined for use in fibre photometry. *Agrp-ires-cre* mice were obtained from Jackson Laboratory *Agrp^tm1(cre)Low/J^* (stock no. 012899) and bred with GHSR1a floxed mice provided by Professor Jeffrey Zigman from the University of Texas SouthWestern at Dallas [52] to delete GHSR from AgRP neurons. For in vivo photometry studies, *Agrp^cre/wt^::Ghsr^wt/wt^* mice were used as control animals (designated as AgRP GHSR WT) and A*grp^cre/wt^::Ghsr^lox/lox^* mice were used as experimental mice (designated as AgRP GHSR KO) to allow for cre-dependent expression of GCaMP6s specifically in AgRP neurons. For food intake and anxiety-like behavioural studies, *Agrp^wt/wt^:: Ghsr^lox/lox^* mice were used as control animals (designated as AgRP GHSR WT) and A*grp^cre/wt^::Ghsr^lox/lox^* mice were used as experimental mice (designated as AgRP GHSR KO). *Dat-ires-cre* mice from Jackson Laboratory (B6.SJL-*Slc6a3^tm1.1(cre)Bkmn^*/J; stock no. 006660) were used to examine the effect of peripheral ghrelin administration on dopamine neural activity in the Ventral Tegmental Area (VTA).

### 1.2 Food intake and motivation studies

Home cage Feeding Experimental Devices 3 (FED3) [53] were placed in home cages of individual housed mice. Mice were trained to collected chow diet pellet (Energy [kcal/g] from protein 24.1%, fat 10.4, carbohydrate 65.5; 5TUM, TestDiets, CA, USA) under fixed ratios (FR) of 1, 3 and 5, where mice are trained to nose poke 1, 3 or 5 times to collect a single pellet in FR1, FR3 or FR5 respectively. To measure food intake in response to ghrelin (1mg/kg) or saline, mice were injected intraperitoneally (ip) during the light phase when the natural feeding drive is low and chow pellet consumption was measured on an FR1 schedule and recorded with FED3s for 90 minutes. To measure ghrelin-induced motivation (ip ghrelin 1mg/kg or saline), mice underwent a progressive ratio (PR) schedule session in which the number of nose pokes required to collect a pellet progressive increases with each pellet delivered. The PR session was conducted during the light phase when natural motivation levels are low and recorded with FED3s for 90 minutes,

### 1.3 Stereotaxic Surgery

Stereotaxic surgeries were performed on adult male at least 10 weeks of age. Mice were anaesthetised with 2-3% isoflurane (Baxter, Australia) and injected with Metacam (5mg/kg) prior to placing into a heatpad-mounted (37C) stereotaxic frame (Stoelting). For GCaMP6s expression in AgRP neurons, cre-dependent AAV9-hSyn-FLEX-GCaMP6s-WPRE-SV40 (∼2.0 x 10^12^ vg/ml; Addgene #100845) was bilaterally injected into the ARC (Coordinates: – 1.6 mm anterior-posterior; 0.2 mm lateral; –5.8 mm dorsoventral from the surface of the brain, 200nl/side infused at a rate of 40nl/min and allowed to rest for 5 minutes post-infusion). For experiments using GCaMP6s in *Dat-ires-cre* mice, cre-dependent AAV9-hSyn-FLEX-GCaMP6s-WPRE-SV40 (∼2.0 x 10^12^ vg/ml; Addgene #100845) was injected unilateral into the VTA (Coordinates: –3.1 mm anterior-posterior; 0.5 mm lateral; –4.4 mm dorsoventral from the surface of the brain, 200nl/side infused at a rate of 40nl/min and allowed to rest for 5 minutes post-infusion). For experiments involving dopamine release, mice were unilaterally injected with the non-cre dependent dopamine biosensor – GRAB-DA (∼2.0 x 10^12^ vg/ml, WZ Biosciences, MD, USA; YL10012-AAV9: AAV-hSyn-DA4.3) in the nucleus accumbens (bregma 1.2 mm, 0.5 mm lateral, –4.8 mm from the surface of the brain; 200nl infused at a rate of 40nl/min and allowed to rest for 5 minutes post-infusion). Ferrule-capped fibres (400 μm core, NA 0.48 Doric, MF1.25 400/430–0.48) were placed above the site of injection and secured with dental cement (GBond, Japan).

In an additional experiment, *Agrp^cre/wt^::Ghsr^wt/wt^*and A*grp^cre/wt^::Ghsr^lox/lox^* mice were injected with both cre-dependent AAV9-hSyn-FLEX-GCaMP6s-WPRE-SV40 (as above) and cre-dependent AAV5-hSyn-DIO-hM3D(Gq)-mCherry (∼2.0 x 10^12^ vg/ml; Addgene #44361) bilaterally injected into the ARC to test the functional capacity of AgRP GHSR KO to increase food intake to a ghrelin independent signal. Mice were given a recovery period of two weeks after surgery and to allow for viral transduction before experimentation.

### 1.4 Fibre Photometry

All recordings were performed using connectorized LEDs, LED drivers, Fluorescent Mini Cubes, rotary joints and photoreceivers (model no. 2151; Newport) from Doric Lenses (Quebec, Canada). These optical components were controlled by a Tucker Davis Technologies (TDT) RZ5P processor using TDT Synapse software for demodulation, low-pass filtering (4Hz) and data acquisition. Two excitation wavelengths were used to deliver 465 nm and 405 nm, in which the 465 nm wavelength reported a Ca2+ dependent GCaMP6 signal or dopamine-specific GRABDA signal and the 405 nm wavelength served as an isosbestic control for motion artifact. The isosbestic wavelength is where excitation is independent from intracellular Ca2+ (GCaMP6) or extracellular dopamine release (GRABDA).

Behavioural events were marked by the researcher using Synapse during recordings or using Open Scope software (Tucker-Davis Technologies) after recordings to precisely align behavioural events with neural activity data. For data processing, custom written python codes extracted and down sampled 465 nm and 405 nm signals to every 100 ms (codes available at Github). These down sampled 465 nm and 405 nm signals were then used to calculate ΔF/F using the following equation (F465 nm-F405 nm/F405 nm) to correct for photobleaching of the signal and movement artefacts. Photobleaching was negligible due to the short experimental time frame (∼30-45 minutes).

For data analysis, z-score normalisation was used for each behavioural event (ie pellet drop, injection) where the degree of change is relative to a predefined baseline period. Z-score normalisation used the following formula; z = (F-Fµ)/Fσ, where F is the signal and Fµ and Fσ are the mean and standard deviation of the baseline signal. A z-score highlights the number of standard deviations a data point is away from the baseline mean, ie a z-score of 5 is 5 standard deviations away from the baseline mean period.

### 1.5 Fibre photometry behavioural experiments

To measure AgRP neural activity and dopamine release in the NAc, mice with fibre implants were habituated to Reese’s peanut butter chips (Fat 29%, Carbohydrate 52%, Protein 3%) in home cages for 3 days prior to experimental recordings. Mice had either ad libitum access to chow diet (fed state) and were fasted overnight for 14 hours (fasted state). Mice were habituated to the photometry setup prior to experimentation and on the day of experiments, mice had 10 minutes to acclimatize before starting recordings. In two-minute intervals, a small wooden dowel (non food object; chewing control) was dropped into the enclosure followed by a chow pellet and peanut butter (PB) pellets. PB pellets were created by dropping melted Reese’s peanut butter chips onto parafilm with a 10ml syringe, such that all PB pellets were an approximately uniform size (average weighed 50mg). This ensures an assessment of AgRP activity or dopamine release to PB consumption is standardised and any differences are related to genotyping rather than total calories consumed and mice always consumed the PB pellet in all trials. For experiments involving IP injection of saline or ghrelin (1mg/kg; BOC Sciences) or CNO (1/mg/kg; Sigma Aldrich, in saline), baseline GCaMP6s activity was measured for 15 minutes prior to injection, at least 25 minutes after injection without food and for a further 10 minutes after food presentation (GCaMP6s studies). For GRABDA studies, baseline dopamine signal was measured for 15 minutes prior to injection before exposure to PB pellets. All photometry recordings were conducted during the early light phase.

### 1.6 Anxiety-like and exploratory behaviours

Ad libitum or overnight fasted AgRP GHSR WT and KO mice were tested in elevated plus maze (EPM), light dark box (LD box) or open field arenas, as previously described [3]. Each trial was video tracked for and analysed with Ethovision (Noldus Information Technology; NL) to quantify mouse performance within defined regions of behavioural arenas. The order in which the mice were assessed was counterbalanced between each test to minimise order effects. The arenas were thoroughly cleaned with 70% ethanol between each trial. A minimum of 1 week was required between repeated exposures of the same mouse to the same arena. Each test was performed during the early light phase to align with the time of GCaMP6 or GRABDA recordings.

### 1.7 Immunohistochemistry

The efficiency of viral transduction was confirmed by immunohistochemical detection of GFP expression in the ARC of *Agrp^cre/wt^::Ghsr^wt/wt^,* A*grp^cre/wt^::Ghsr^lox/lox^* or the VTA of *Dat^cre/wt^* mice. Mice were perfused and fixed with 0.05M phosphate buffered saline (PBS) followed by 4% paraformaldehyde (PFA) in PBS with 4% paraformaldehyde. Brains were immediately removed and post-fixed for 24 hours in 4% PFA at 4°C before being transferred to 30% sucrose 0.1M PB solution for until the suspended brain sunk in the sucrose solution. Brains were processed coronally in a cryostat (Leica CM1800) at −20°C and were sectioned at 30µm. Sections were collected in sets of 4 and stored in 24-well plates suspended in cryoprotectant at −20°C. Sections were washed in 0.1M PBS (3 x 10m mins) and then blocked for 60 minutes in 4% normal horse serum (NHS) in PBs + 0.3% Triton-X. Primary antibodies (chicken anti-GFP; ab13970; abcam) was added to blocking solution at a dilution of 1:1000 and incubated overnight at 4^0^C on an orbital shaker. The next day, sections were washed in 0.1M PB (3 x 10 mins) and incubated with 488 secondary antibody (goat anti-chicken; AB_2337390; Jackson ImmunoResearch) at a dilution of 1:500 in 0.1M PB for 2 hours at room temperature. Following a final series of washing to remove the secondary antibody secondary antibody, sections were mounted onto SuperFrost slides (Thermo Fisher Scientific) with VECTASHIELD antifade mounting medium with DAPI (Vector Labs). Post-curing, slides were then sealed with nail polish and stored at 4°C. Sections were imaged under an upright fluorescence microscope (Zeiss Axio Imager 2; Zeiss, Germany). Coronal brain slices were outlined and tiled with an EC Plan-NEOFLUAR 5x objective (0,15NA, air), while focal images were captured through an EC Plan-NEOFLUAR 10x objective (0,3NA, air). DAPI and 488 staining were imaged in combination with BFP and GFP filters respectively paired with a high-power mercury arc lamp. Prior to tiling acquisition, exposure, gain and offset were adjusted to the brightest spot on the sample. Care was taken to ensure no pixels were oversaturated.

### 1.8 Statistical Analysis

Data are represented as mean ± SEM and all statistical analyses were performed using GraphPad Prism for MacOS X. Two-way ANOVAs (with repeated measures as appropriate) and post hoc tests were used to determine statistical significance. A two-tailed Student’s t-test was used when appropriate, as indicated in figure legends. p < 0.05 was considered statistically significant.

## Results

### 3.1 AgRP^GHSRs^ regulate food intake, motivation and AgRP neuronal activity

To demonstrate the functional effects of GHSR deletion in AgRP neurons, we examined ghrelin-induced chow pellet intake and motivation using FED3 operant devices (Fig 1A)[53]. Although ip ghrelin (1mg/kg) significantly increased chow consumption over 90 mins in a FR1 feeding schedule in AgRP GHSR WT mice, it failed to induce chow pellet intake in AgRP GHSR KO mice (Fig 1B). Furthermore, AgRP neurons and ip ghrelin are both stimulate motivated food seeking using PR schedules [16; 54], therefore we injected ip ghrelin (1mg/kg) to test whether AgRP^GHSRs^ are necessary for motivated chow food seeking using a PR. IP ghrelin increased the number of pellets consumed during a PR schedule in WT but not KO mice (Fig 1C), with a significant difference between pellets collected in AgRP GHSR WT and KO mice after ip ghrelin injection. To show that AgRP^GHSR^ are required for ghrelin to increase AgRP neuronal activity, we used an in vivo photometry approach in AgRP GHST WT and KO mice. IP ghrelin injections rapidly increased AgRP neuronal activity in WT but not KO mice (main effect of genotype), with significant differences in averaged Z-scores at 5-minute time bins after injection (Fig 1H, 1I). The subsequent response to chow consumption 20 mins after ghrelin injection cause a greater suppression of AgRP neuronal activity from 0-5 minutes in WT compared to KO mice and no significant difference was observed from 5-10 minutes (Fig 1J, 1K). This coincided with greater chow consumption in AgRP GHSR WT mice during the recording period (Fig 1L). No genotype differences in AgRP activity were observed in response to IP saline and chow presentation (Fig 1M-P), although chow presentation suppressed AgRP activity (main effect of time; Fig 1P). Furthermore, co-transduction of AgRP neurons with hM3Gq DREADD and GCaMP6s revealed that CNO equally stimulated AgRP activity in both AgRP GHSR WT and KO mice (main effect of time; Sup Fig 1), indicating that AgRP neurons respond normally to other stimuli in the absence of GHSR expression. Collectively, these studies illustrate that our genetic approach to delete GHSR in AgRP neurons prevents ghrelin-induced food intake, motivation and AgRP neuronal activity without perturbing neuronal integrity, confirming that this is a useful model for further studies.

**Figure 1:**
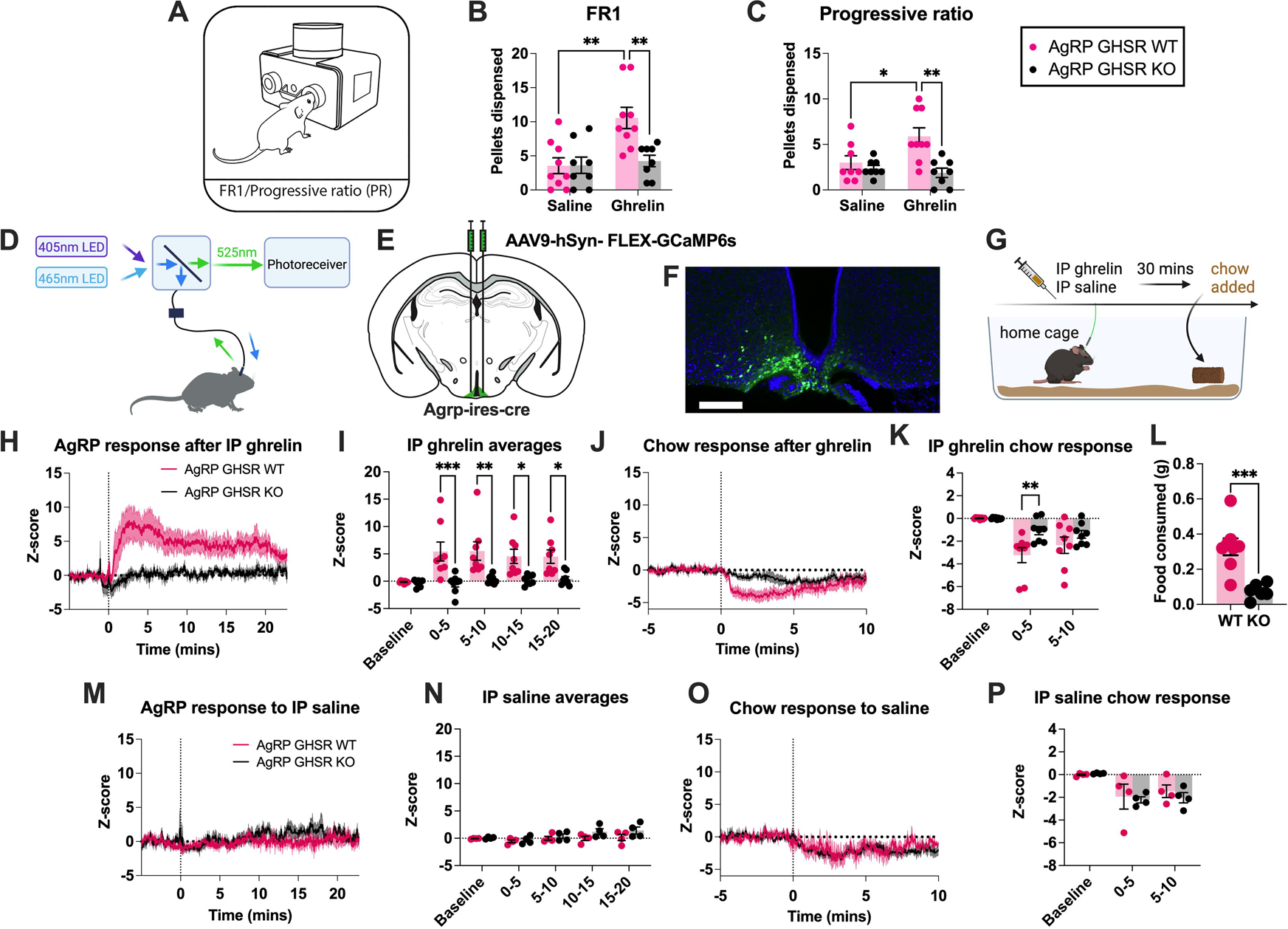
Deletion of GHSR in AgRP neurons affects food intake, motivation and AgRP neuronal activity. (A) Schematic representation of a FED3 device used for home cage assessment of food intake, via a fixed ratio 1 (FR1) schedule (B; WT n=9; KO n=8), or food motivation using a progressive ratio (PR) schedule (C; WT n=8; KO n=8). (D) Schematic representation of a photometry setup, with GCaMP6s excitation at 465 nm and an isosbestic control (405nm). (E) Schematic of AAV9-hSyn-FLEX-GCaMP6s injection into the ARC of AgRP cre mice. (F) GCaMP6s expression in the ARC of AgRP cre mice; scale bar 100µm. (G) Schematic experimental timeline of photometry experiments created with BioRender.com. The averaged Z-score of AgRP neuronal responses to IP ghrelin aligned to injection at time = 0 sec (H, WT n=8; KO n=8) and averaged Z-score time bins between 0-5, 5-10, 10-15 and 15-20 minutes (I; WT n=8; KO n=8). AgRP neuronal response to chow consumption in AgRP GHSR WT and KO mice (J; WT n=8; KO n=8) with average Z-score time bins showing a greater fall in AgRP activity in WT, compared to KO mice (K). 1hr food intake measured during the IP ghrelin photometry recordings in WT and KO mice (L; n=8 WT and KO). The averaged Z-score response (M), or the time binned responses (N), to IP saline was not different in WT compared to KO mice (n=4). In response to chow diet consumption, there was no difference in the averaged Z-score over time (O) or in 5-minute time bins (P). Data +/− SEM. Dotted lines in H and M represent the time of injection and in J & O represent time chow was placed into the cage. Two-way ANOVA with post hoc Sidak’s multiple comparisons (B, C, I, K) or students t-test (L). * p<0.05, ** p<0.01, *** p<0.001. For a detailed description of statistics see statistical table 1.

### 3.2 AgRP^GHSRs^ influence AgRP neural activity in response to sensory information

Sensory detection of food and/or food-specific cues rapidly inhibit AgRP neurons in a manner that is scaled to the metabolic need of the animal and caloric availability of the food or food predicting cue [27; 28; 32]. This highlights that AgRP neurons rely on both internal and external information to correctly respond to food and/or cues. While calorie consumption provides AgRP neurons the opportunity to update and encode calorie value with external sensory cues via gut-brain feedback [28; 30; 31], exactly how internal cues of metabolic need integrate with external sensory cues to influence AgRP activity remains unknown. To test if GHSRs on AgRP neurons influence AgRP activity in response to external sensory information, we expressed GCaMP6s in both AgRP GHSR WT and KO mice. Small equally-sized PB pellets were used as a high-caloric value food item and a wooden dowel was used as a chewing control item. In the fed state, the introduction of a wooden dowel resulted in a difference in AgRP activity between genotype (main effect of genotype), although no specific differences were identified by post hoc analysis (Fig 2C). The presentation of PB pellets to ad libitum fed mice significantly affected AgRP activity (main effect of time) and genotype (main effect of genotype), with post hoc analysis indicating a trend for an attenuated suppression in average Z-score for 0-60 secs in KO compared to WT mice (Fig 2E).

**Figure 2:**
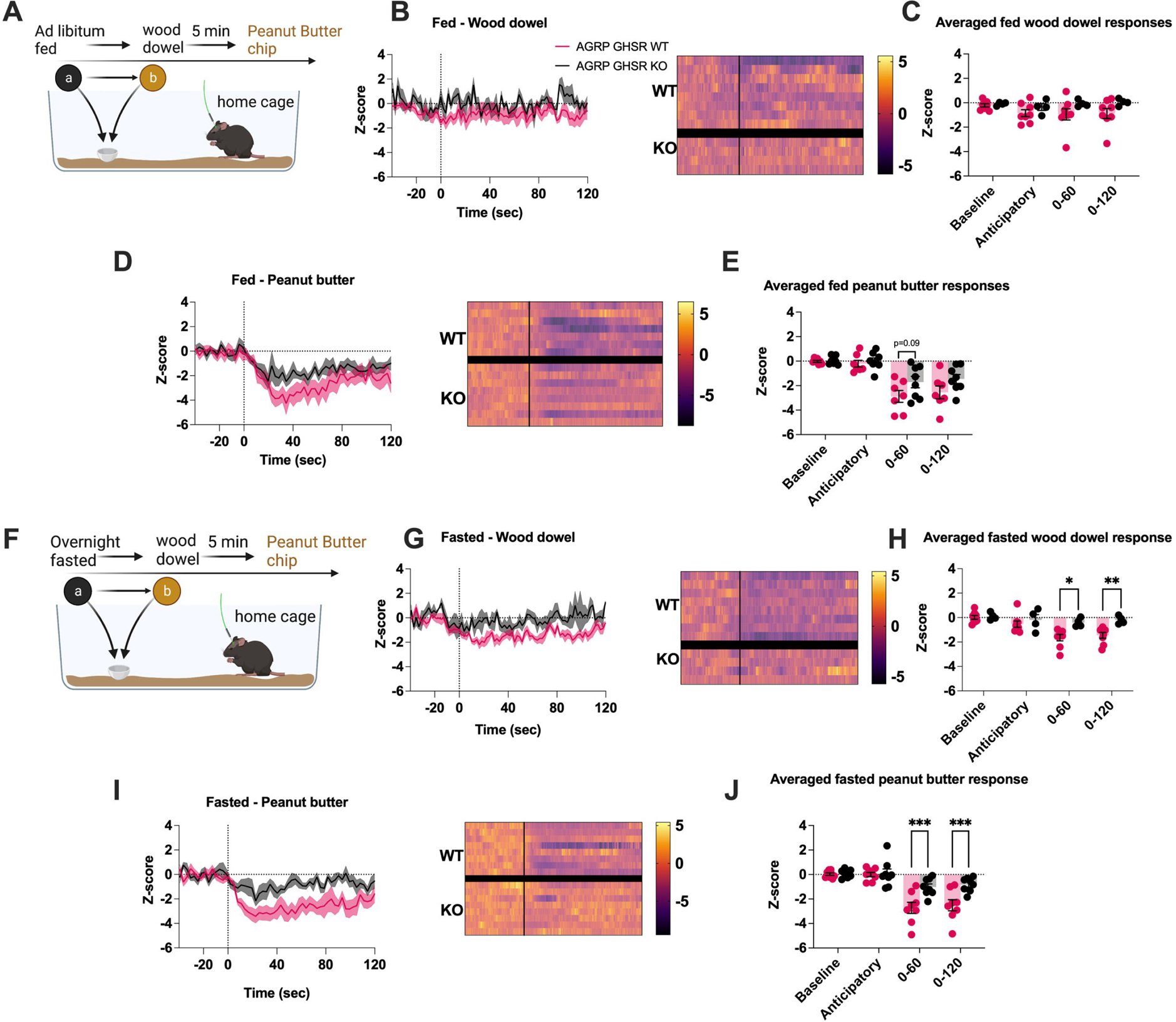
Deletion of GHSR in AgRP neurons attenuates fasting-induced response to a wood dowel and peanut butter chips. Assessment of AgRP neuronal activity in ad-libitum fed AgRP GHSR WT or KO mice to a wooden dowel and peanut butter pellet (PB) as schematically represented (A; created with BioRender.com). Average Z-score traces of AgRP neuronal activity in response to a wooden dowel aligned to first contact of the nose to the object with heat maps (B; WT n=8, KO n=4) and the following time averaged bins in baseline (−40 to −20 sec), anticipatory (−20 to 0 sec), the first minute (0 to 60 secs) and 2 minutes (0-120 secs) (C; WT n=8, KO n=4). Average Z-score traces in response to peanut butter aligned to first contact with heat maps (D) and the following time averaged bins in baseline (−40 to −20 sec), anticipatory (−20 to 0 sec), the first minute (0 to 60 secs) and 2 minutes (0-120 secs) (E; WT n=7, KO n=8). Assessment of AgRP neuronal activity in overnight fasted AgRP GHSR WT or KO mice to a wooden dowel and peanut butter as schematically represented (F; created with BioRender.com). Average Z-score traces in response to a wooden dowel aligned to first contact of the nose to the object and heat maps (G) and the following time averaged bins in baseline (−40 to −20 sec), anticipatory (−20 to 0 sec), the first minute (0 to 60 secs) and 2 minutes (0-120 secs) (H; WT n=8, KO n=4). Average Z-score traces in response to peanut butter aligned to first contact (I) and the following time averaged bins in baseline (−40 to −20 sec), anticipatory (−20 to 0 sec), the first minute (0 to 60 secs) and 2 minutes (0-120 secs) (J). Data +/− SEM. Dotted lines in B, D, G, I represent first contact. Two-way ANOVA with post hoc Sidak’s multiple comparisons (H, J). * p<0.05, ** p<0.01, *** p<0.001. For a detailed description of statistics see statistical table 1.

In fasted mice, the introduction of a wooden dowel resulted in lower AgRP activity (main effect of time), which was attenuated in KO mice across all time bins examined (main effect of genotype). Moreover, post hoc analysis revealed significantly less suppression of AgRP activity in KO mice at 0-60 and 0-120 minutes after presentation of a wooden dowel (Fig 2H). The presentation of equally-sized PB pellets, to ensure equally caloric consumption across genotypes and trials, resulted in a difference in AgRP activity over time (main effect of time), genotype (main effect of genotype) and difference between genotypes over time (interaction time x genotype; Fig 2J). Post hoc analysis demonstrated the fall in AgRP activity in KO mice at 0-60 secs and 0-120 secs was significantly attenuated compared to WT mice (Fig 2J). Importantly, no differences in anxiety-like or exploratory behaviour were detected between AgRP GHSR WT and KO mice in EPM, LD box or Open Field tests under ad libitum fed or fasted metabolic states (Sup Fig 2 and Sup Fig 3). These data suggest that the attenuated fall in AgRP activity to PB pellets or wooden dowel in KO mice was not caused by altered anxiety-like or exploratory behaviour, an important consideration as both AgRP neurons and the ghrelin system play a role in stress and anxiety-like behaviour [42-44; 47; 48].

### 3.3 AgRP^GHSRs^ influence ghrelin-induced NAc dopamine release

Previous studies highlight that AgRP neurons influence the development of dopamine neuroplasticity [55] and dopamine release in the NAc [45]. Indeed, impaired metabolic sensing in AgRP neurons reduces dopamine-driven motivation during fasting and reduces striatal dopamine release in response to palatable foods or during operant sucrose seeking [32]. Thus, we used GRAB-DA sensors to assess the impact of sensory cues (wooden dowel, PB pellets) on dopamine release in response to ip saline and ghrelin in AgRP GHSR WT and AgRP GHSR KO mice. An ip injection of saline had no effect on NAc dopamine release in WT or KO mice (Fig 3C), however, ip ghrelin resulted in a significant difference in dopamine release over time (main effect of time) and between genotypes (main effect of genotype). Post hoc analysis indicating dopamine release was significantly lower average Z-score in KO mice 15-30 secs after dowel presentation (Fig 3E). PB pellet presentation after ip saline results in a significant difference over time (main effect of time) and between genotypes (main effect of genotype; Fig 3G), although no specific differences were identified by post hoc analysis (Fig 3G). In contrast, NAc dopamine release after ip ghrelin was significantly reduced in AgRP GHSR KO compared to WT mice at 0-15 secs after PB pellet presentation, as assessed by post hoc analysis (Fig 3I).

**Figure 3:**
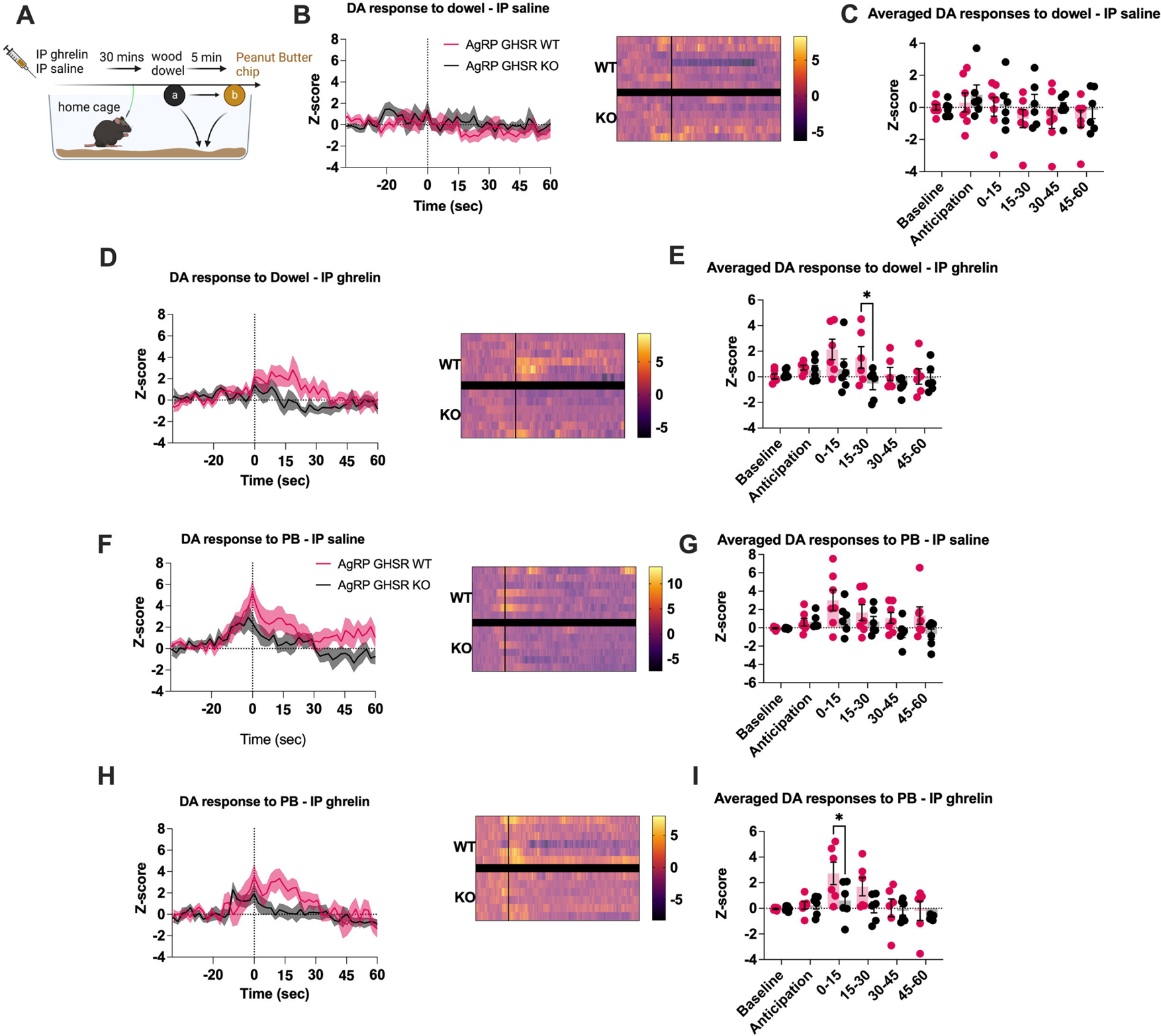
Deletion of GHSR in AgRP neurons affects ghrelin-induced changes in dopamine (DA) release in the nucleus accumbens. Schematic of the experimental timeline created with BioRender.com (A), where mice expressing GRAB_DA_ in the nucleus accumbens were injected with either ghrelin or saline and presented with a wooden dowel followed by a peanut butter chip (PB). Averaged Z-score traces of dopamine release in the NAc in response to a wooden dowel following IP ghrelin with heat maps (B; WT n=7, KO n=6) or saline with heat maps (D; WT n=7, KO n=6) and respective time binned averages (C and E) organised into baseline (−40 to −20 sec), anticipatory (−20 to 0 sec) periods and 15-second periods after PB/dowel (0-15, 15-30, 30-45, 45-60). Averaged Z-score traces of dopamine release to PB following IP ghrelin (F WT n=7, KO n=6) or saline (H; WT n=6, KO n=6) with their respective time bin averages (G and I). Data +/− SEM. Dotted lines in B, D, F, H represent first contact to wood dowel or PB. Two-way ANOVA with post hoc Sidak’s multiple comparisons (E, I). * p<0.05. For a detailed description of statistics see statistical table 1.

### 3.4 IP ghrelin does not direct regulate VTA dopamine neurons in vivo

A large number of studies suggest that ghrelin increases motivation and dopamine release by targeting the VTA [37; 39; 56-59], however the reduced dopamine release after ip ghrelin to both wooden dowel and PB pellets in AgRP GHSR KO suggests a novel role for GHSR signalling in AgRP neurons. To explore how ip ghrelin influences VTA DA neural activity, we used in vivo photometry with GCaMP6s injection into the VTA of DAT-ires-cre mice (Fig 4A). Surprisingly, ip ghrelin injection had no effect on the population activity of VTA dopamine neurons (Fig 4B-C). However, ip ghrelin injection acutely increased the population activity of VTA DA neurons to subsequent presentation of PB pellets of chow 10-20 minutes after injection (Fig 4D-G). These studies suggest that peripheral ghrelin does not directly regulate VTA dopamine neural activity but rather acts on upstream neural circuits that provide input to the VTA.

**Figure 4:**
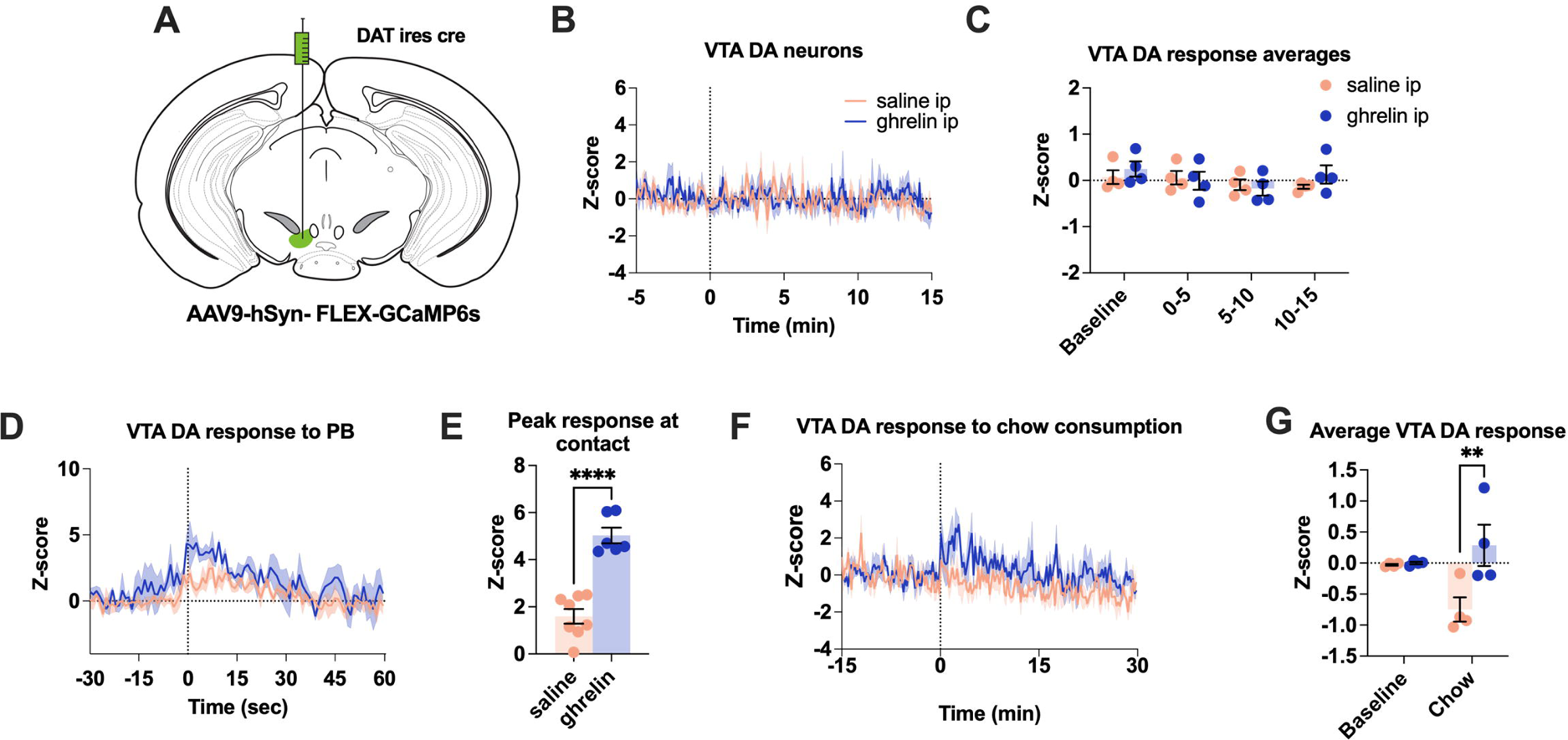
Ghrelin-induced changes in VTA dopamine neurons. Schematic showing the approach to record from DA neurons after injection of AAV9-hSyn-FLEX-GCaMP6s in the ventral tegmental area (VTA) in dopamine transporter (DAT) ires cre mice (A). Average Z-score VTA DA response to an injection of ghrelin or saline (B; saline & ghrelin n=4; dotted line represents time at injection) with 5 minute time binned data (C). VTA dopamine neuronal activity at contact with PB (D; saline n=4, ghrelin n=3), with the maximum Z-score response at contact to after saline or ghrelin injection (E, saline n= 8 contacts from n=4 mice; ghrelin n=6 contacts from n=3 mice). (F) Average Z-score VTA DA response to chow consumption (saline n=3, ghrelin n=4; dotted represents time when chow was placed in cage). (G) The averaged 30-minute Z-score response from the beginning of chow consumption after saline or ghrelin injection (saline n= 3; ghrelin n=4). Data +/− SEM. Dotted lines in B, D, F represent first contact to PB or chow. Two-way ANOVA with post hoc Sidak’s multiple comparisons (H, J). * p<0.05, ** p<0.01, *** p<0.001, **** p<0.0001. For a detailed description of statistics see statistical table 1.

### 3.5 AgRP^GHSRs^ do not affect fasting-induced NAc dopamine release

Next we assessed whether GHSR signalling in AgRP neurons influences NAc dopamine to sensory cues (wooden dowel and PB pellets) under fasted conditions, since fasting and energy deficit is a strong driver of ghrelin secretion into the bloodstream [35]. The presentation of a wooden dowel (Fig 5C) or PB pellets (Fig 5E) to ad libitum fed WT and KO mice affected dopamine release over time (main effect of time) although no significant difference between genotypes were observed (Fig 5C, 5E). Similarly, a wooden dowel or PB pellet affected dopamine release over time (main effect of time; Fig 5G, 5I) with no significant difference between genotypes (Fig 5G, 5I). These studies highlight that AgRP^GHSRs^ do not affect NAc dopamine release in response to fasting.

**Figure 5:**
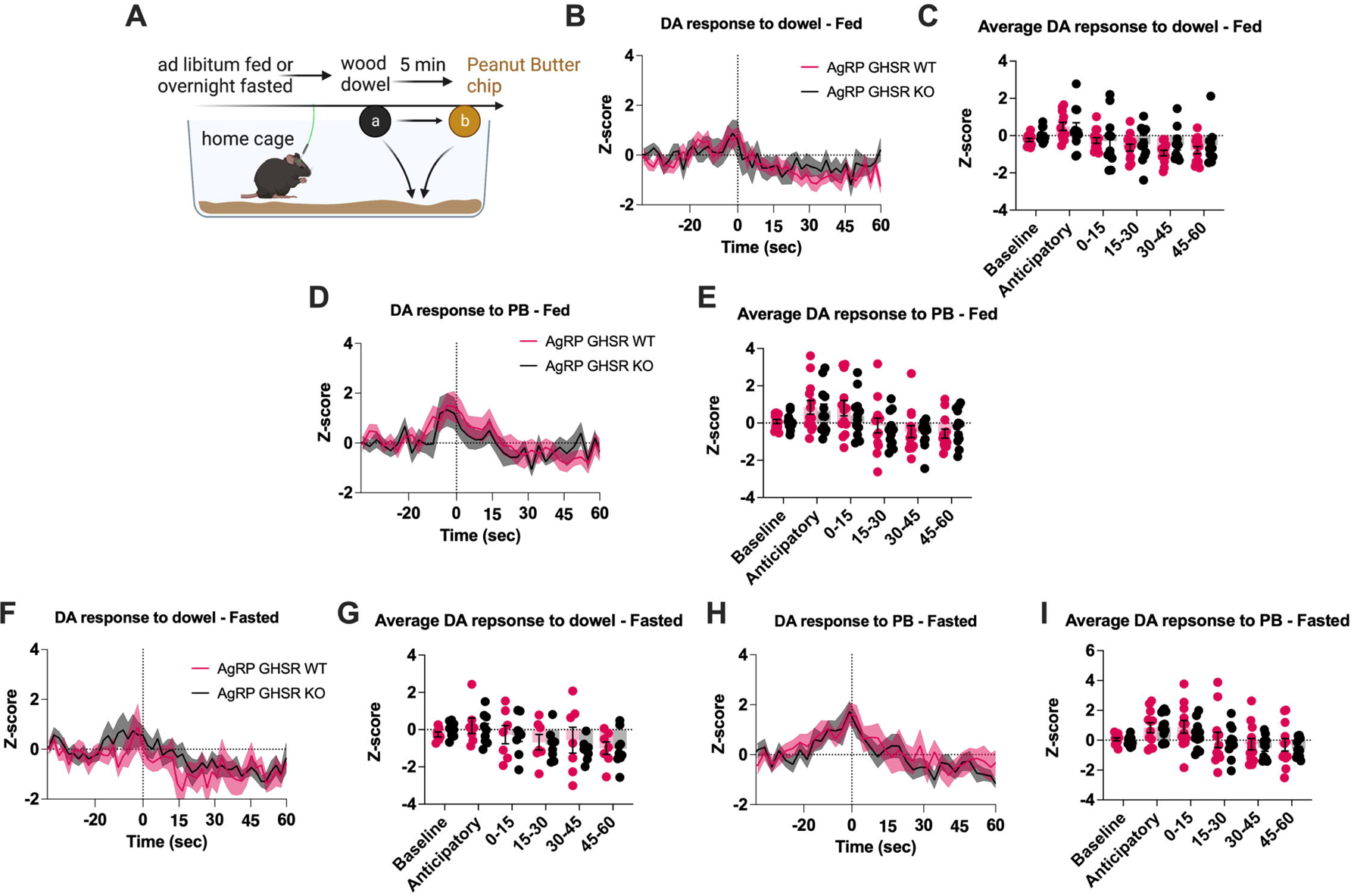
Fasting did not reveal GHSR-mediated NAc dopamine release. (A) Schematic of experimental timeline created with BioRender.com where mice expressing GRAB-DA had ad-libitum access to chow or were fasted overnight before being introduced to a wooden dowel and peanut butter. Averaged Z-score traces in ad libitum fed mice aligned with point of first contact with a wooden dowel (B; WT n=12, KO n=13) and respective time averaged bins (C; WT n=12, KO n=13). Averaged Z-score traces aligned with point of first contact with peanut butter (D; WT n=12, KO n=13) and respective time averaged bins (E; WT n=12, KO n=13). Averaged Z-score traces in fasted mice aligned with point of first contact with a wooden dowel (F) and respective time averaged bins (G). Averaged Z-score traces with peanut butter (H) and respective time averaged bins (I). Data +/− SEM. Dotted lines in B, D, F, H represent first contact to wood dowel or PB. For a detailed description of statistics see statistical table 1.

## 4.1 Discussion

Recent studies show that the sensory detection of caloric availability suppresses AgRP neuronal activity prior to meaningful food consumption in a manner that is dependent on interoceptive energy need of the animal. While manipulating the caloric value and availability of external food or food cues clearly demonstrates an effect on AgRP neuronal activity [9; 26-28; 32], how interoceptive signals of energy need influence AgRP activity in response to the sensory detection of food or food cues remains unexplored. In this study, we explored whether AgRP^GHSRs^ integrated interoceptive energy need with the sensory detection of food (PB) and a non-food object (wooden dowel). We found that deleting GHSRs from AgRP neurons impaired ghrelin-induced induced chow intake, food motivation and AgRP neuronal activity, confirming the functional GHSR deletion in AgRP neurons KO mice. These results are supported by studies showing that AgRP neurons are an important target for plasma ghrelin. For example, the majority of AgRP neurons contain GHSRs [60] and AgRP neurons are largely responsible for the ability of ghrelin and fasting to increase food intake and affect energy expenditure [50; 61-64]. Further, food deprivation-induced discrimination studies [6; 36] show AgRP neurons sense hunger whereas plasma ghrelin signals hunger. Thus, GHSR receptors on AgRP neurons are a critical component of the negative feedback actions of ghrelin to hunger.

However, our novel results show that AgRP^GHSRs^ influence the response of AgRP neurons to external sensory cues, suggesting ghrelin signaling on AgRP neurons is more than just a feedback signal to increase AgRP activity during hunger. Importantly, the suppression in AgRP activity after the introduction of sensory cues (food – PB, object - wooden dowel) was significantly attenuated in fasted, but not fed, AgRP GHSR KO mice. These results are in line with previous studies demonstrating that AgRP inhibition to food requires the inherent ability of AgRP neurons to metabolise and sense changes in glucose [32]. Thus, our results suggest that AgRP^GHSRs^ are required to integrate the interoceptive energy state with external sensory information to produce the optimal response in AgRP neural activity.

We have recently suggested AgRP neurons regulate energy balance through a process of energy allostasis, rather than negative feedback to perturbations in homeostasis [4]. Energy allostasis incorporates previous experiences to help predict and prepare for perceived future energy demands prior to an energy deficit and if an energy deficit does occur, homeostatic systems work to restore energy balance. For example, sensory cues cause a greater suppression of AgRP neural activity, prior to food consumption, when these cues were previously associated with calorie consumption [28; 34]. Intriguingly, AgRP^GHSRs^ influence both the fall in AgRP activity to the sensory detection of food or food cues and act as hunger signals to increase AgRP activity in response to energy need. Therefore, we suggest AgRP^GHSRs^ maintain energy balance through energy allostasis rather than a negative feedback model of energy homeostasis.

Of note, AgRP neuronal responses to a non-food object (a wooden dowel) were also significantly lower in a manner that depended on GHSR expression in AgRP neurons. Although AgRP neurons are classically studied for the role in energy homeostasis and feeding behaviour, recent studies show that sensory information of non-food related events also rapidly suppress AgRP neuronal activity, including thermal pain, maternal reunion, warm exposure (14C→30C) and cessation of running [12; 65-67]. The functional consequences of rapid AgRP inhibition to the sensory detection of non-food related stimuli is unknown, yet our results show it is dependent on metabolic state and AgRP^GHSR^ expression. We hypothesize this may be related to an important role in foraging behaviour since the need to forage is greater when hungry and a foraging individual is likely to encounter both food and non-food related objects or events. Indeed, AgRP neurons and ghrelin play important roles in foraging and foraging-related behaviours such as locomotion, exploration and arousal ([68–71] and AgRP neurons alter behaviours based on energy need [72].

We noted that ghrelin did not increase motivation in AgRP GHSR KO mice. These results are consistent with the idea that AgRP neurons increase food motivation and food seeking [16; 32; 73], but they highlight an essential role for AgRP^GHSRs^ for ghrelin-induced motivation. Although ghrelin is well described to influence food reward and motivation, these effects have been ascribed to actions within the mesolimbic dopamine system based on direct brain injections of ghrelin into the VTA or pharmacological and genetics approaches targeting VTA neural activity [37; 39; 56; 58; 59; 74]. Thus, our results highlight a new pathway for ghrelin to influence motivation by acting on AgRP^GHSRs^.

The presence of GHSRs on AgRP neurons is also important for normal dopamine release in the NAc in response to the introduction of a wood dowel or PB pellet. AgRP GHSR KO mice had an attenuated response to PB at 0-15 secs and dowel 15-30 secs after presentation. While previous studies show that ip ghrelin injections increases dopamine release or turnover in the NAc [58; 75], the exact site of action was unknown. Our studies directly implicate a role for AgRP^GHSR^ to mediate ghrelin-induced dopamine release. Indeed at important role for AgRP neurons is supported by the literature since AgRP play an important role in motivated feeding behaviour and engage midbrain dopaminergic neurons in the ventral tegmental area (VTA), as well as influencing dopamine release in the nucleus accumbens (NAc) [45; 76]. Recently, we showed that metabolic sensing in AgRP neurons was required to translate internal energy need into increased motivated behaviour [32]. A metabolic-sensing impairment in AgRP neurons also reduced NAc dopamine release to food rewards or during motivated food seeking and attenuated the ability to learn the caloric value of peanut butter [32]. These studies highlight that AgRP neurons actively transmit ghrelin as an interoceptive signal of hunger to influence NAc dopamine release. However, it should be noted that no significant differences in NAc dopamine release between WT and KO mice were observed in overnight fasted mice, suggesting other pathways must also be involved in fasted-induced dopamine release, such as the intrinsic metabolic-sensing ability of AgRP neurons [32]. Similar to AgRP activity, GHSR deletion in AgRP neurons also attenuated dopamine release after wooden dowel and PB pellet presentation in response to ghrelin injection. These results suggest that ghrelin-signaling in AgRP neurons affects the salience of both food and non-food events through changes in NAc dopamine release. Again, this may be related to optimal foraging behaviour since foraging involves the exploration and interaction with food and non-food objects/events and involves dopamine release in the NAc [77; 78].

Interestingly, we did not observe a direct effect of ghrelin injection on VTA dopamine population activity although ghrelin potentiated dopamine neural activity to PB or chow presentation. The lack of a direct effect on VTA dopamine activity is somewhat surprising giving the abundant expression of GHSRs on dopamine neurons [79] and well known role of intra VTA ghrelin injection on food intake and behaviour [37; 39; 57; 59; 80]. The potentiated VTA dopamine activity to PB or chow presentation after ghrelin injection suggests an important role of indirect ghrelin sensitive pathway. While the nature of this input is unknown, AgRP^GHSR^ neurons are a strong possibility given the ability to AgRP neurons to influence VTA dopamine activity [76] and the attenuated dopamine release to ghrelin in AgRP GHSR KO mice in the current study. Other possibilities include an interaction with orexin neurons in the lateral hypothalamus since the effects of ghrelin on food reward are absent in orexin KO mice [38].

In summary, our results demonstrate that AgRP^GHSRs^ influence both AgRP neural activity and NAc dopamine release to the sensory detection of food and non-food objects. While ghrelin was always considered an important feedback signal to defend against weight loss and starvation [81], these results highlight the novel possibility that ghrelin signalling in AgRP neurons affects AgRP neuronal activity to the sensory detection of food and food cues. Thus, AgRP^GHSRs^ integrate an interoceptive energy state with current external sensory information to produce an optimal response in AgRP neural activity. In this manner, ghrelin signaling in AgRP neurons controls energy balance through a process of energy allostasis[4], in which the integration of energy need with the current sensory information of caloric information is likely to facilitate optimal behaviour when exposed to similar sensory cues in the future. This may be a novel principle in which neural hunger-sensing accelerates learning.

## Supporting information

Supplemental Statistical Table 1

## Acknowledgements

This study was supported by an NHMRC grant and fellowship to ZBA (1126724, 1154974). We acknowledge that BioRender was used to produce elements incorporated in the figure and graphical abstract (Biorender.com).

---

**Supplementary figure 1: Deletion of GHSR from AgRP neurons does not affect DREADD stimulation of AgRP neurons.** AgRP cre mice were co-injected with AAV9-hSyn-FLEX-GCaMP6s and AAV5-hSyn-DIO-hM3D(Gq)-mCherry to transduce AgRP neurons with both GCAmP6s and hM3D(Gq). CNO increased average Z-score over time (A) and averaged time binned Z-scores (B) in both AGRP GHSR WT (n=9) and KO mice (n=8). Data +/− SEM. Dotted lines in A represents time at injection of CNO. Two-way ANOVA main effect of time (B). For a detailed description of statistics see statistical table 1.

**Supplement Figure 2: Deletion of GHSRs in AgRP neurons does not affect anxiety-like behaviour in ad libitum fed mice.** Anxiety-like behaviour in ad libitum fed AgRP GHSR WT or AGRP GHSR KO mice was assessed using Elevated Plus Maze (EPM), Light Dark Box (LD Box) and Open Field test. In the EPM, no differences in distance moved (A), number of open arm entries (B) or time in the open arm (C) were observed. There were no differences in distance moved (D), number of light zone entries (E) or time in the light zone (F) in the LD Box test. Similar for the Open Field test, there were no differences in distance moved (G), number of inner zone entries (H) or time in the inner zone (I). Data +/− SEM; AgRP GHSR WT n=10, AgRP GHSR KO n=11; student’s t-test. For a detailed description of statistics see statistical table 1.

**Supplement Figure 3: Deletion of GHSRs in AgRP neurons does not affect anxiety-like behaviour in overnight fasted mice.** The Elevated Plus Maze (EPM), Light Dark Box (LD Box) and Open Field test were used to examine anxiety-like behaviour in overnight fasted (14-16 hrs) AgRP GHSR WT or AgRP GHSR KO mice. There were no differences in distance moved (A), number of open arm entries (B) or time in the open arm (C) of the EPM. In the LD Box, no differences in distance moved (D), number of light zone entries (E) or time in the light zone (F) were observed. Similar for the Open Field test, there were no differences in distance moved (G), number of inner zone entries (H) or time in the inner zone (I). Data +/− SEM; AgRP GHSR WT n=10, AgRP GHSR KO n=11; student’s t-test. For a detailed description of statistics see statistical table 1.

## References

[1] Caron, A., Richard, D., 2017. Neuronal systems and circuits involved in the control of food intake and adaptive thermogenesis. Ann N Y Acad Sci 1391(1):35–53.

[2] Ferrario, C.R., Labouebe, G., Liu, S., Nieh, E.H., Routh, V.H., Xu, S., et al., 2016. Homeostasis Meets Motivation in the Battle to Control Food Intake. J Neurosci 36(45):11469–11481.

[3] Clarke, R.E., Voigt, K., Reichenbach, A., Stark, R., Bharania, U., Dempsey, H., et al., 2023. Identification of a Stress-Sensitive Anorexigenic Neurocircuit From Medial Prefrontal Cortex to Lateral Hypothalamus. Biol Psychiatry 93(4):309–321.

[4] Reed, F., Lockie, S.H., Reichenbach, A., Foldi, C.J., Andrews, Z.B., 2022. Appetite to learn: An allostatic role for AgRP neurons in the maintenance of energy balance. Current Opinion in Endocrine and Metabolic Research 24(June 2022).

[5] Sutton, A.K., Krashes, M.J., 2020. Integrating Hunger with Rival Motivations. Trends in endocrinology and metabolism: TEM 31(7):495–507.

[6] Siemian, J.N., Arenivar, M.A., Sarsfield, S., Aponte, Y., 2021. Hypothalamic control of interoceptive hunger. Current Biology 31:1–13.

[7] Jennings, J.H., Ung, R.L., Resendez, S.L., Stamatakis, A.M., Taylor, J.G., Huang, J., et al., 2015. Visualizing hypothalamic network dynamics for appetitive and consummatory behaviors. Cell 160(3):516–527.

[8] Hahn, T.M., Breininger, J.F., Baskin, D.G., Schwartz, M.W., 1998. Coexpression of Agrp and NPY in fasting-activated hypothalamic neurons. Nat Neurosci 1(4):271–272.

[9] Mandelblat-Cerf, Y., Ramesh, R.N., Burgess, C.R., Patella, P., Yang, Z.F., Lowell, B.B., et al., 2015. Arcuate hypothalamic AgRP and putative POMC neurons show opposite changes in spiking across multiple timescales. Elife 4.

[10] Murphy, B.A., Fioramonti, X., Jochnowitz, N., Fakira, K., Gagen, K., Contie, S., et al., 2009. Fasting enhances the response of arcuate neuropeptide Y-glucose-inhibited neurons to decreased extracellular glucose. Am J Physiol Cell Physiol 296(4):C746–756.

[11] Liu, T.M., Kong, D., Shah, B.P., Ye, C.P., Koda, S., Saunders, A., et al., 2012. Fasting Activation of AgRP Neurons Requires NMDA Receptors and Involves Spinogenesis and Increased Excitatory Tone. Neuron 73(3):511–522.

[12] Deem, J.D., Faber, C.L., Pedersen, C., Phan, B.A., Larsen, S.A., Ogimoto, K., et al., 2020. Cold-induced hyperphagia requires AgRP neuron activation in mice. Elife 9.

[13] Aponte, Y., Atasoy, D., Sternson, S.M., 2011. AGRP neurons are sufficient to orchestrate feeding behavior rapidly and without training. Nature Neuroscience 14(3):351–355.

[14] Cavalcanti-de-Albuquerque, J.P., Bober, J., Zimmer, M.R., Dietrich, M.O., 2019. Regulation of substrate utilization and adiposity by Agrp neurons. Nat Commun 10.

[15] Chen, Y.M., Lin, Y.C., Zimmerman, C.A., Essner, R.A., Knight, Z.A., 2016. Hunger neurons drive feeding through a sustained, positive reinforcement signal. Elife 5.

[16] Krashes, M.J., Koda, S., Ye, C., Rogan, S.C., Adams, A.C., Cusher, D.S., et al., 2011. Rapid, reversible activation of AgRP neurons drives feeding behavior in mice. J Clin Invest 121(4):1424–1428.

[17] Bewick, G.A., Gardiner, J.V., Dhillo, W.S., Kent, A.S., White, N.E., Webster, Z., et al., 2005. Post-embryonic ablation of AgRP neurons in mice leads to a lean, hypophagic phenotype. FASEB J 19(12):1680–1682.

[18] Gropp, E., Shanabrough, M., Borok, E., Xu, A.W., Janoschek, R., Buch, T., et al., 2005. Agouti-related peptide-expressing neurons are mandatory for feeding. Nature Neuroscience 8(10):1289–1291.

[19] Luquet, S., Perez, F.A., Hnasko, T.S., Palmiter, R.D., 2005. NPY/AgRP neurons are essential for feeding in adult mice but can be ablated in neonates. Science 310(5748):683–685.

[20] Morrison, C.D., Morton, G.J., Niswender, K.D., Gelling, R.W., Schwartz, M.W., 2005. Leptin inhibits hypothalamic Npy and Agrp gene expression via a mechanism that requires phosphatidylinositol 3-OH-kinase signaling. Am J Physiol Endocrinol Metab 289(6):E1051–1057.

[21] Schwartz, M.W., Sipols, A.J., Marks, J.L., Sanacora, G., White, J.D., Scheurink, A., et al., 1992. Inhibition of hypothalamic neuropeptide Y gene expression by insulin. Endocrinology 130(6):3608–3616.

[22] Andrews, Z.B., Liu, Z.W., Walllingford, N., Erion, D.M., Borok, E., Friedman, J.M., et al., 2008. UCP2 mediates ghrelin’s action on NPY/AgRP neurons by lowering free radicals. Nature 454(7206):846–851.

[23] Briggs, D.I., Andrews, Z.B., 2011. Metabolic status regulates ghrelin function on energy homeostasis. Neuroendocrinology 93(1):48–57.

[24] Baver, S.B., Hope, K., Guyot, S., Bjorbaek, C., Kaczorowski, C., O’Connell, K.M., 2014. Leptin modulates the intrinsic excitability of AgRP/NPY neurons in the arcuate nucleus of the hypothalamus. J Neurosci 34(16):5486–5496.

[25] Huang, Y., He, Z., Gao, Y., Lieu, L., Yao, T., Sun, J., et al., 2018. Phosphoinositide 3-Kinase Is Integral for the Acute Activity of Leptin and Insulin in Male Arcuate NPY/AgRP Neurons. J Endocr Soc 2(6):518–532.

[26] Betley, J.N., Xu, S., Cao, Z.F., Gong, R., Magnus, C.J., Yu, Y., et al., 2015. Neurons for hunger and thirst transmit a negative-valence teaching signal. Nature 521(7551):180–185.

[27] Chen, Y., Lin, Y.C., Kuo, T.W., Knight, Z.A., 2015. Sensory detection of food rapidly modulates arcuate feeding circuits. Cell 160(5):829–841.

[28] Su, Z.W., Alhadeff, A.L., Betley, J.N., 2017. Nutritive, Post-ingestive Signals Are the Primary Regulators of AgRP Neuron Activity. Cell Rep 21(10):2724–2736.

[29] Garfield, A.S., Shah, B.P., Burgess, C.R., Li, M.M., Li, C., Steger, J.S., et al., 2016. Dynamic GABAergic afferent modulation of AgRP neurons. Nature Neuroscience 19(12):1628–1635.

[30] Beutler, L.R., Chen, Y.M., Ahn, J.S., Lin, Y.C., Essner, R.A., Knight, Z.A., 2017. Dynamics of Gut-Brain Communication Underlying Hunger. Neuron 96(2):461-+.

[31] Goldstein, N., McKnight, A.D., Carty, J.R.E., Arnold, M., Betley, J.N., Alhadeff, A.L., 2021. Hypothalamic detection of macronutrients via multiple gut-brain pathways. Cell Metabolism 33:1–12.

[32] Reichenbach, A., Clarke, R.E., Stark, R., Lockie, S.H., Mequinion, M., Dempsey, H., et al., 2022. Metabolic sensing in AgRP neurons integrates homeostatic state with dopamine signalling in the striatum. Elife 11.

[33] Jikomes, N., Ramesh, R.N., Mandelblat-Cerf, Y., Andermann, M.L., 2016. Preemptive Stimulation of AgRP Neurons in Fed Mice Enables Conditioned Food Seeking under Threat. Current Biology 26(18):2500–2507.

[34] Berrios, J., Li, C., Madara, J.C., Garfield, A.S., Steger, J.S., Krashes, M.J., et al., 2021. Food cue regulation of AGRP hunger neurons guides learning. Nature 595(7869):695-+.

[35] Zigman, J.M., Bouret, S.G., Andrews, Z.B., 2016. Obesity Impairs the Action of the Neuroendocrine Ghrelin System. Trends in Endocrinology and Metabolism 27(1):54–63.

[36] Davidson, T.L., Kanoski, S.E., Tracy, A.L., Walls, E.K., Clegg, D., Benoit, S.C., 2005. The interoceptive cue properties of ghrelin generalize to cues produced by food deprivation. Peptides 26(9):1602–1610.

[37] Abizaid, A., Liu, Z.W., Andrews, Z.B., Shanabrough, M., Borok, E., Elsworth, J.D., et al., 2006. Ghrelin modulates the activity and synaptic input organization of midbrain dopamine neurons while promoting appetite. J Clin Invest 116(12):3229–3239.

[38] Perello, M., Sakata, I., Birnbaum, S., Chuang, J.C., Osborne-Lawrence, S., Rovinsky, S.A., et al., 2010. Ghrelin increases the rewarding value of high-fat diet in an orexin-dependent manner. Biol Psychiatry 67(9):880–886.

[39] Skibicka, K.P., Hansson, C., Alvarez-Crespo, M., Friberg, P.A., Dickson, S.L., 2011. Ghrelin directly targets the ventral tegmental area to increase food motivation. Neuroscience 180:129–137.

[40] Theander-Carrillo, C., Wiedmer, P., Cettour-Rose, P., Nogueiras, R., Perez-Tilve, D., Pfluger, P., et al., 2006. Ghrelin action in the brain controls adipocyte metabolism. J Clin Invest 116(7):1983–1993.

[41] Wortley, K.E., Anderson, K.D., Garcia, K., Murray, J.D., Malinova, L., Liu, R., et al., 2004. Genetic deletion of ghrelin does not decrease food intake but influences metabolic fuel preference. Proc Natl Acad Sci U S A 101(21):8227–8232.

[42] Lutter, M., Sakata, I., Osborne-Lawrence, S., Rovinsky, S.A., Anderson, J.G., Jung, S., et al., 2008. The orexigenic hormone ghrelin defends against depressive symptoms of chronic stress. Nat Neurosci 11(7):752–753.

[43] Spencer, S.J., Xu, L., Clarke, M.A., Lemus, M., Reichenbach, A., Geenen, B., et al., 2012. Ghrelin regulates the hypothalamic-pituitary-adrenal axis and restricts anxiety after acute stress. Biol Psychiatry 72(6):457–465.

[44] Stark, R., Santos, V.V., Geenen, B., Cabral, A., Dinan, T., Bayliss, J.A., et al., 2016. Des-Acyl Ghrelin and Ghrelin O-Acyltransferase Regulate Hypothalamic-Pituitary-Adrenal Axis Activation and Anxiety in Response to Acute Stress. Endocrinology 157(10):3946–3957.

[45] Alhadeff, A.L., Goldstein, N., Park, O., Klima, M.L., Vargas, A., Betley, J.N., 2019. Natural and Drug Rewards Engage Distinct Pathways that Converge on Coordinated Hypothalamic and Reward Circuits. Neuron 103(5):891-+.

[46] Burnett, C.J., Li, C., Webber, E., Tsaousidou, E., Xue, Stephen Y., Brüning, Jens C., et al., 2016. Hunger-Driven Motivational State Competition. Neuron 92(1):187–201.

[47] Dietrich, M.O., Zimmer, M.R., Bober, J., Horvath, T.L., 2015. Hypothalamic Agrp Neurons Drive Stereotypic Behaviors beyond Feeding. Cell 160(6):1222–1232.

[48] Wang, C.M., Zhou, W.J., He, Y., Yang, T., Xu, P.W., Yang, Y.J., et al., 2021. AgRP neurons trigger long-term potentiation and facilitate food seeking. Transl Psychiatry 11(1).

[49] Lee, J.H., Xue, B., Chen, Z., Sun, Y., 2022. Neuronal GHS-R Differentially Modulates Feeding Patterns under Normal and Obesogenic Conditions. Biomolecules 12(2).

[50] Wang, Q., Liu, C., Uchida, A., Chuang, J.C., Walker, A., Liu, T., et al., 2014. Arcuate AgRP neurons mediate orexigenic and glucoregulatory actions of ghrelin. Mol Metab 3(1):64–72.

[51] Wu, C.S., Bongmba, O.Y.N., Yue, J., Lee, J.H., Lin, L., Saito, K., et al., 2017. Suppression of GHS-R in AgRP Neurons Mitigates Diet-Induced Obesity by Activating Thermogenesis. Int J Mol Sci 18(4).

[52] Gupta, D., Patterson, A.M., Osborne-Lawrence, S., Bookout, A.L., Varshney, S., Shankar, K., et al., 2021. Disrupting the ghrelin-growth hormone axis limits ghrelin’s orexigenic but not glucoregulatory actions. Mol Metab 53:101258.

[53] Matikainen-Ankney, B.A., Earnest, T., Ali, M., Casey, E., Sutton, A.K., Legaria, A., et al., 2021. Feeding Experimentation Device version 3 (FED3): An open-source home-cage compatible device for measuring food intake and operant behavior. Elife 10.7554/eLife.66173.

[54] Finger, B.C., Dinan, T.G., Cryan, J.F., 2012. Diet-induced obesity blunts the behavioural effects of ghrelin: studies in a mouse-progressive ratio task. Psychopharmacology (Berl) 220(1):173–181.

[55] Dietrich, M.O., Bober, J., Ferreira, J.G., Tellez, L.A., Mineur, Y.S., Souza, D.O., et al., 2012. AgRP neurons regulate development of dopamine neuronal plasticity and nonfood-associated behaviors. Nat Neurosci 15(8):1108–1110.

[56] Jerlhag, E., Egecioglu, E., Dickson, S.L., Andersson, M., Svensson, L., Engel, J.A., 2006. Ghrelin stimulates locomotor activity and accumbal dopamine-overflow via central cholinergic systems in mice: implications for its involvement in brain reward. Addict Biol 11(1):45–54.

[57] Jerlhag, E., Egecioglu, E., Dickson, S.L., Douhan, A., Svensson, L., Engel, J.A., 2007. Ghrelin administration into tegmental areas stimulates locomotor activity and increases extracellular concentration of dopamine in the nucleus accumbens. Addict Biol 12(1):6–16.

[58] Jerlhag, E., Egecioglu, E., Dickson, S.L., Engel, J.A., 2010. Glutamatergic regulation of ghrelin-induced activation of the mesolimbic dopamine system. Addict Biol Jun 23 [Epub ahead of print].

[59] Skibicka, K.P., Shirazi, R.H., Rabasa-Papio, C., Alvarez-Crespo, M., Neuber, C., Vogel, H., et al., 2013. Divergent circuitry underlying food reward and intake effects of ghrelin: dopaminergic VTA-accumbens projection mediates ghrelin’s effect on food reward but not food intake. Neuropharmacology 73:274–283.

[60] Willesen, M.G., Kristensen, P., Romer, J., 1999. Co-localization of growth hormone secretagogue receptor and NPY mRNA in the arcuate nucleus of the rat. Neuroendocrinology 70(5):306–316.

[61] Chen, H.Y., Trumbauer, M.E., Chen, A.S., Weingarth, D.T., Adams, J.R., Frazier, E.G., et al., 2004. Orexigenic action of peripheral ghrelin is mediated by neuropeptide Y and agouti-related protein. Endocrinology 145(6):2607–2612.

[62] Luquet, S., Phillips, C.T., Palmiter, R.D., 2007. NPY/AgRP neurons are not essential for feeding responses to glucoprivation. Peptides 28(2):214–225.

[63] Mani, B.K., Osborne-Lawrence, S., Mequinion, M., Lawrence, S., Gautron, L., Andrews, Z.B., et al., 2017. The role of ghrelin-responsive mediobasal hypothalamic neurons in mediating feeding responses to fasting. Mol Metab 6(8):882–896.

[64] Wu, C.S., Bongmba, O.Y.N., Yue, J., Lee, J.H., Lin, L.G., Saito, K., et al., 2017. Suppression of GHS-R in AgRP Neurons Mitigates Diet-Induced Obesity by Activating Thermogenesis. International Journal of Molecular Sciences 18(4).

[65] Alhadeff, A.L., Su, Z., Hernandez, E., Klima, M.L., Phillips, S.Z., Holland, R.A., et al., 2018. A Neural Circuit for the Suppression of Pain by a Competing Need State. Cell 173(1):140–152 e115.

[66] Miletta, M.C., Iyilikci, O., Shanabrough, M., Sestan-Pesa, M., Cammisa, A., Zeiss, C.J., et al., 2020. AgRP neurons control compulsive exercise and survival in an activity-based anorexia model. Nature Metabolism 2(11):1204–1211.

[67] Zimmer, M.R., Fonseca, A.H.O., Iyilikci, O., Pra, R.D., Dietrich, M.O., 2019. Functional Ontogeny of Hypothalamic Agrp Neurons in Neonatal Mouse Behaviors. Cell 178(1):44–59 e47.

[68] Keen-Rhinehart, E., Bartness, T.J., 2005. Peripheral ghrelin injections stimulate food intake, foraging, and food hoarding in Siberian hamsters. American journal of physiology. Regulatory, integrative and comparative physiology 288(3):R716–722.

[69] Vestlund, J., Bergquist, F., Eckernas, D., Licheri, V., Adermark, L., Jerlhag, E., 2019. Ghrelin signalling within the rat nucleus accumbens and skilled reach foraging. Psychoneuroendocrinology 106:183–194.

[70] Huang, A., Maier, M.T., Vagena, E., Xu, A.W., 2023. Modulation of foraging-like behaviors by cholesterol-FGF19 axis. Cell Biosci 13(1):20.

[71] Liu, Q., Yang, X., Luo, M., Su, J., Zhong, J., Li, X., et al., 2023. An iterative neural processing sequence orchestrates feeding. Neuron 111(10):1651–1665 e1655.

[72] Burnett, C.J., Funderburk, S.C., Navarrete, J., Sabol, A., Liang-Guallpa, J., Desrochers, T., et al., 2019. Need-based prioritization of behavior. Elife 8.

[73] Chen, Y.M., Essner, R.A., Kosar, S., Miller, O.H., Lin, Y.C., Mesgarzadeh, S., et al., 2019. Sustained NPY signaling enables AgRP neurons to drive feeding. Elife 8.

[74] Chuang, J.C., Perello, M., Sakata, I., Osborne-Lawrence, S., Savitt, J.M., Lutter, M., et al., 2011. Ghrelin mediates stress-induced food-reward behavior in mice. The Journal of clinical investigation 121(7):2684–2692.

[75] Jerlhag, E., 2008. Systemic administration of ghrelin induces conditioned place preference and stimulates accumbal dopamine. Addict Biol 13(3-4):358–363.

[76] Mazzone, C.M., Liang-Guallpa, J., Li, C., Wolcott, N.S., Boone, M.H., Southern, M., et al., 2020. High-fat food biases hypothalamic and mesolimbic expression of consummatory drives. Nature Neuroscience 23(10):1253-+.

[77] Floresco, S.B., Seamans, J.K., Phillips, A.G., 1996. A selective role for dopamine in the nucleus accumbens of the rat in random foraging but not delayed spatial win-shift-based foraging. Behav Brain Res 80(1-2):161–168.

[78] Hills, T.T., 2006. Animal foraging and the evolution of goal-directed cognition. Cogn Sci 30(1):3–41.

[79] Zigman, J.M., Jones, J.E., Lee, C.E., Saper, C.B., Elmquist, J.K., 2006. Expression of ghrelin receptor mRNA in the rat and the mouse brain. J Comp Neurol 494(3):528–548.

[80] Lockie, S.H., Stark, R., Spanswick, D.C., Andrews, Z.B., 2019. Glucose availability regulates ghrelin-induced food intake in the Ventral Tegmental Area. Journal of neuroendocrinology:e12696.

[81] Mani, B.K., Zigman, J.M., 2017. Ghrelin as a Survival Hormone. Trends in Endocrinology and Metabolism 28(12):843–854.

