## Supplementary figures and images for "Ghrelin signalling in AgRP neurons links metabolic state to the sensory regulation of AgRP neural activity"

### Supplemental Figure 1

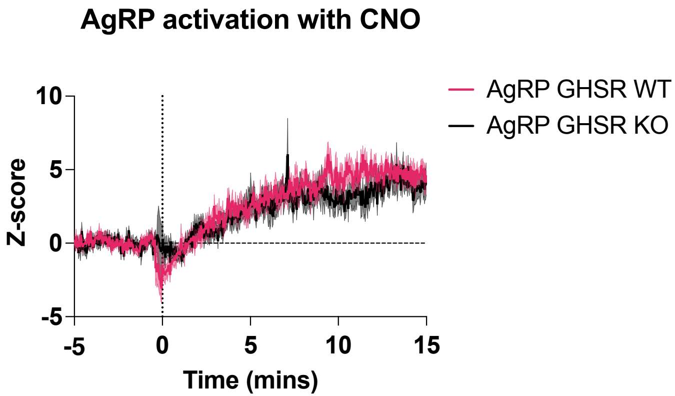

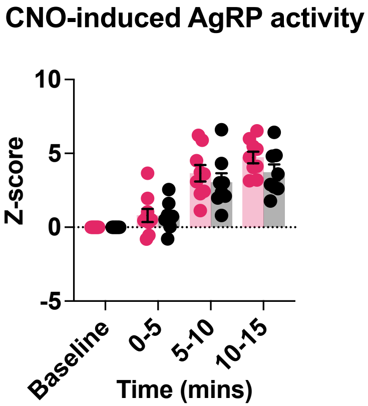

### Supplemental Figure 2

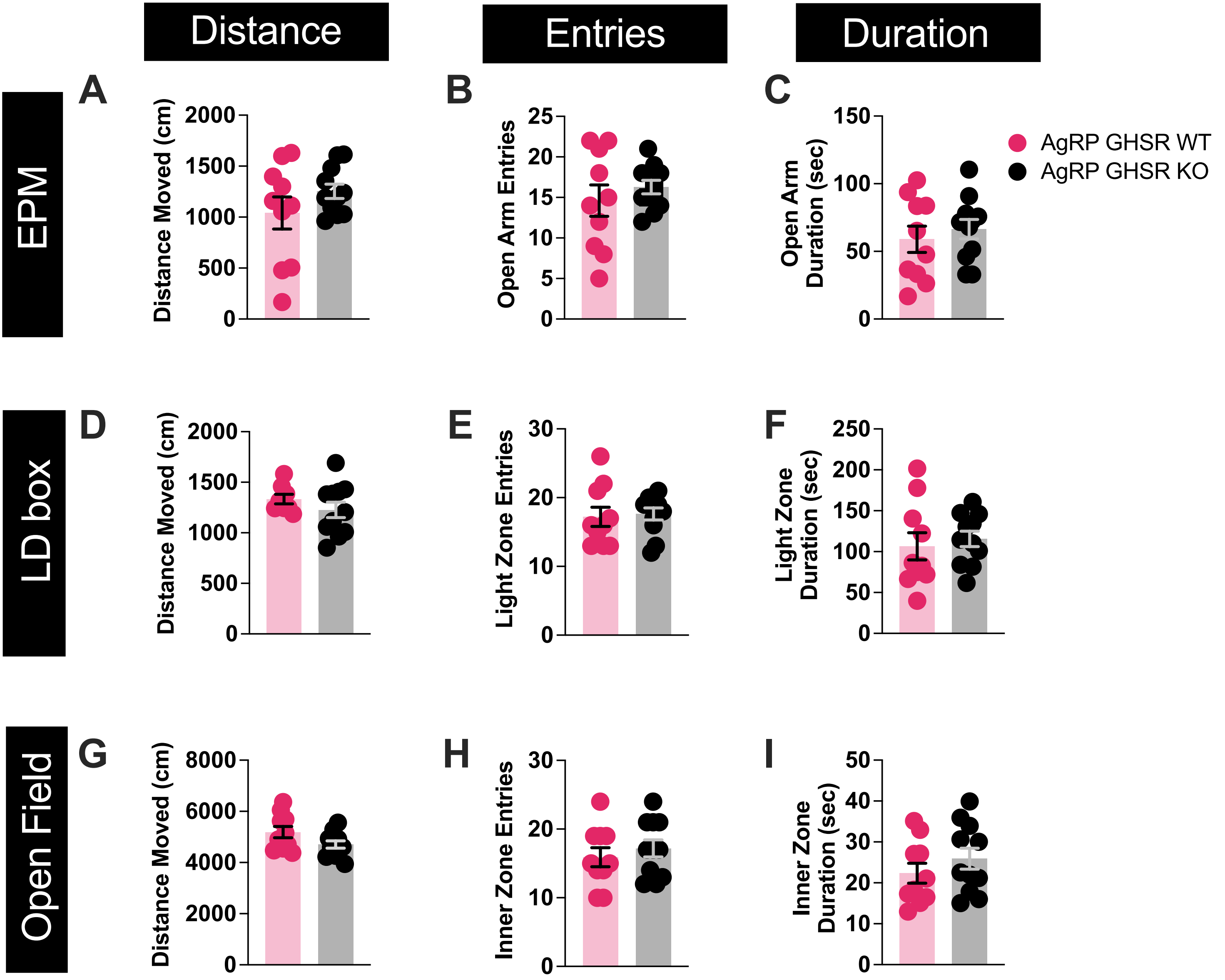

### Supplemental Figure 3

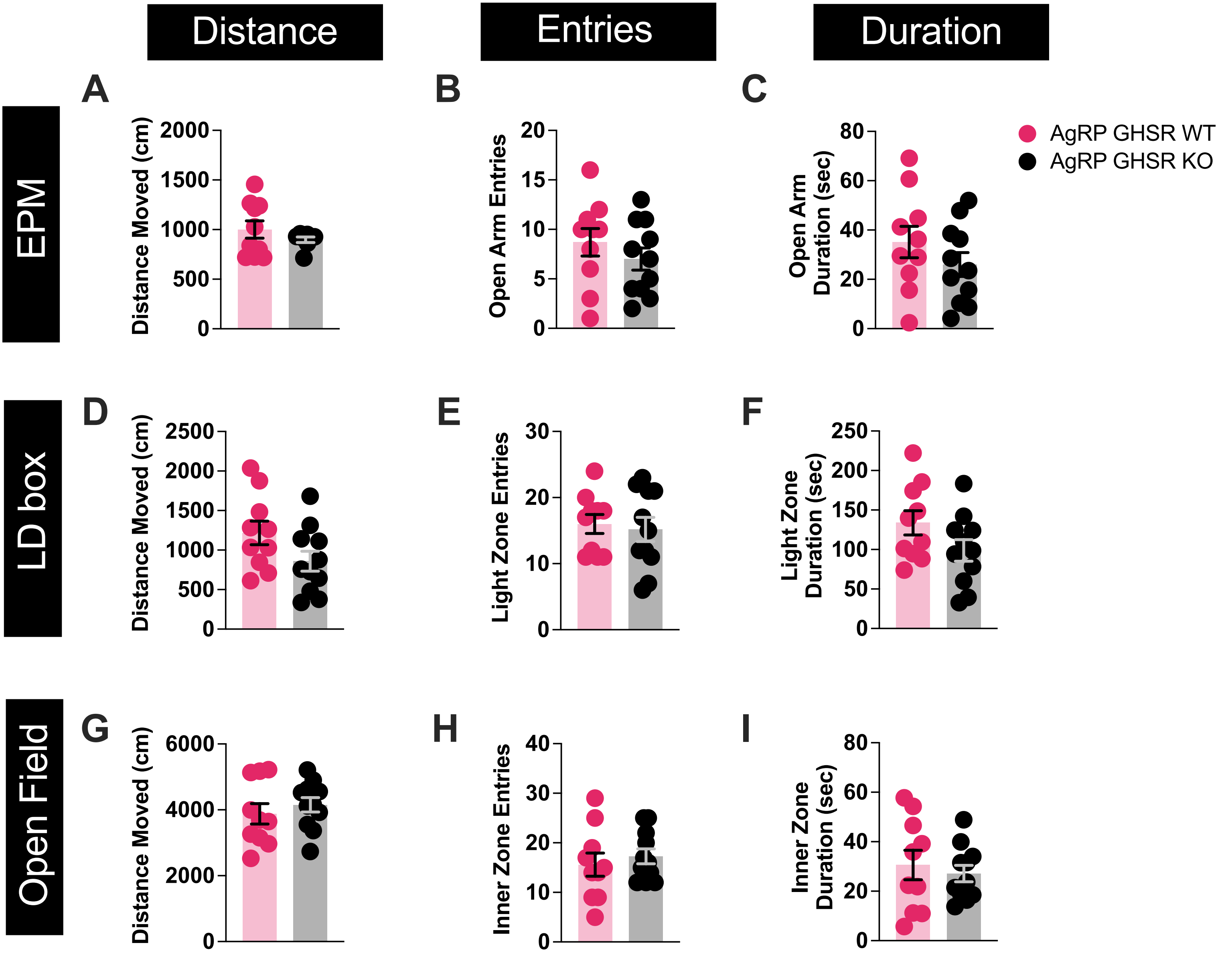
